# Therapeutic effects of Chlorzoxazone, a BKCa channel agonist, in a mouse model of Fragile X syndrome

**DOI:** 10.1101/2020.12.11.389569

**Authors:** Valerie Lemaire-Mayo, Marion Piquemal, Wim E. Crusio, Eric Louette, Susanna Pietropaolo

## Abstract

Fragile X syndrome (FXS) is an X-linked developmental disorder characterized by several behavioral abnormalities, including hyperactivity, sensory hyper-responsiveness and cognitive deficits, as well as autistic symptoms, e.g., reduced social interaction. These behavioural alterations are recapitulated by the major animal model of FXS, i.e., the Fmr1-KO mouse, which has been extensively employed to identify therapeutic targets for FXS, though effective pharmacological treatments are still lacking. Here we focused on the therapeutic role of large-conductance Calcium-dependent potassium (BKCa) channels, playing a crucial role in neuronal excitability and neurotransmitter release. Reduced expression/functionality of these channels has been described in FXS patients and mice, so that molecules activating these channels have been proposed as promising treatments for this syndrome. Here we performed an extensive characterization of the therapeutic impact of a novel BKCa agonist on FXS-like symptoms in the Fmr1-KO mouse model, employing a drug repurposing setting. We evaluated the acute and chronic effects of chlorzoxazone, i.e., a classical drug used for non-developmental muscular pathologies, on the locomotor, social, cognitive and sensory-motor alterations of Fmr1-KO mice and compared them with other pharmacological treatments recently proposed for FXS that instead do not target BKCa channels. Our results clearly demonstrate for the first time the marked efficacy of chlorzoxazone in treating all the behavioral abnormalities of FXS mice, thus encouraging the preferential use of this molecule over others for clinical applications in the field of FXS, and potentially of other neurodevelopmental disorders.

## INTRODUCTION

Fragile X Syndrome (FXS) is the most common form of inherited intellectual disability and the leading monogenic cause of Autism Spectrum Disorder (ASD). It is caused by a mutation in the Fragile X Mental Retardation 1 (*FMR1*) gene on the X chromosome, resulting in a deficiency of Fragile X Mental Retardation Protein, FMRP (Pieretti *et al.*, 1991), an RNA-binding protein that regulates dendritic branching, synaptic function (Santoro *et al.*, 2012), and neuronal maturation (Greenough *et al.*, 2001). The absence of FMRP induces physical anomalies (e.g., facial elongation, macro-orchidism) and several behavioural abnormalities, including hyperactivity, sensory hyper-responsiveness, and cognitive deficits, often associated with typical autistic symptoms such as impaired social interactions (Tranfaglia, 2011).

FXS is successfully modelled by the Fmr1-KO mouse line, recapitulating most of the physical and behavioral abnormalities observed in FXS patients (Jung *et al.*, 2014, Oddi *et al.*, 2013, Pietropaolo *et al.*, 2014, Pietropaolo *et al.*, 2011). In the original Fmr1-knockout (*Fmr1^tm1Cgr^*) mouse, FMRP expression was abrogated by insertional inactivation of exon 5 of the *Fmr1* gene, i.e., the mouse analogue of the human *FMR1* (Dutch-Belgian Fragile X Consortium, 1994). Since the Fmr1 promoter is nonetheless intact, transcription remains possible, resulting in abnormal residual Fmr1 RNAs (Mientjes *et al.*, 2006, Yan *et al.*, 2004). Hence, a complete Fmr1 null mouse was then developed, characterized by the absence of all Fmr1 mRNA transcripts (Mientjes *et al.*, 2006). This FXS second generation model — known as Fmr1-KO2 (Mientjes *et al.*, 2006) — has been largely used for brain studies (De Esch *et al.*, 2015, De Vrij *et al.*, 2008, El Fatimy *et al.*, 2012, Jung *et al.*, 2012, Levenga *et al.*, 2009, Mao *et al.*, 2013, Pilpel *et al.*, 2009, Scotto-Lomassese *et al.*, 2011, Vinueza Veloz *et al.*, 2012, Wijetunge *et al.*, 2014, Zhang *et al.*, 2014), but its behavioural FXS-like phenotype seems less robust than the one of the Fmr1-KO model of first generation (Gaudissard *et al.*, 2017). Surprisingly, only few studies have included a behavioral analysis of the Fmr1-KO2 mouse line (De Vrij *et al.*, 2008, Gantois *et al.*, 2017, Mao *et al.*, 2013, Roberts *et al.*, 2007, Yan *et al.*, 2004).

Indeed, the original Fmr1-KO mouse line (or first generation model) has been widely used to identify therapeutic approaches for FXS (reviewed in (Pietropaolo & Subashi, 2014)). However, to date, there is no effective treatment for this syndrome, as most pharmacological approaches target only few symptoms (e.g., anxiolytics and antipsychotics for irritability and hyperactivity). These treatments not only have limited efficacy, but also have serious side effects that are particularly undesirable due to the developmental nature of this syndrome (Pietropaolo *et al.*, 2017). New therapies are therefore urgently needed for FXS; recently, novel molecules have been proposed based on encouraging results on Fmr1-KO mice, including Metformin (Gantois *et al.*, 2017), a first-line therapy for type 2 diabetes acting on matrix metallo-proteinase 9 (MMP9), and Gaboxadol, a GABA(A) receptor agonist (Cogram *et al.*, 2019, Olmos-Serrano *et al.*, 2011). Nonetheless, the efficacy of these treatments has not been convincingly demonstrated on multiple behavioural deficits and following both acute and chronic administration.

Recent studies have suggested the relevance of Ca2+-activated high-conductance potassium (BKCa) channels as emerging therapeutic targets for neurodevelopmental disorders (NDDs) and FXS in particular (Liang *et al.*, 2019). These channels, activated by membrane depolarization and increased intracellular Ca2+ concentration, regulate neurotransmitter release and neuronal triggering (Sah & Faber, 2002), and modulate the activity of dendrites as well as astrocytes and microglia (Hu *et al.*, 2001). In addition to their role in the central nervous system, BKCa channels are also involved in smooth muscle contraction, endocrine cell secretion and proliferation (Jackson, 2005, Tykocki *et al.*, 2017). They have been used to date as pharmacological targets for the treatment of several medical conditions, including stroke, ischemia, muscular and urinary disorders (Jensen, 2002). The relevance of BKCa channels in the NDDs has emerged only recently (Liang *et al.*, 2019): disrupted expression and functionality of BKCA has been described in a subject with autism (Laumonnier *et al.*, 2006), while a mutation in the *CRBN* gene, a regulator upstream of the BK channel, has been associated with non-syndromic recessive mental retardation (Higgins *et al.*, 2008). Furthermore, we demonstrated reduced expression and functionality of BKCa channels in FXS patients (Hebert *et al.*, 2014) and Fmr1-KO mice (Zhang *et al.*, 2014), supporting their role as therapeutic targets (Carreno-Munoz *et al.*, 2018).

Here we propose a BKCa agonist (Liu *et al.*, 2003), chlorzoxazone (5-chloro-2,3-dihydro-1,3-benzoxazol-2-one; CHLOR), as a novel treatment for FXS. CHLOR is an FDA-approved classical drug, centrally acting as a muscle relaxant (Hohmann *et al.*, 2019), by exerting an effect primarily at the level of the spinal cord and subcortical areas of the brain (Martindale, 1993). Hence, CHLOR already has European marketing authorization and could therefore be rapidly tested in clinical trials using a drug repurposing setting. Recently, CHLOR has emerged as a brand-new value in healthcare, exerting immunomodulatory (Deng *et al.*, 2020) and neuroprotective effects in pre-clinical mouse studies (Bai & Ma, 2020). The present study provides with an exhaustive characterization of the effects of acute and chronic systemic administration of CHLOR on multiple FXS-like behaviours in the Fmr1-KO mouse model, including the evaluation of dose-dependent effects, a replication in the Fmr1-KO2 model of second generation, and a comparison of the efficacy of CHLOR with other proposed treatments for FXS not targeting BKCa channels, i.e., Metformin and Gaboxadol.

## METHODS

### ANIMALS

A total of 6 studies were performed, employing different batches of mice. As commonly done in research studies on mouse models of FXS, only adult males were used, since they are known to exhibit the most robust FXS- and ASD-like behavioural phenotypes, as only limited behavioral deficits have been reported in adult female Fmr1-KOs and younger male mutants (Gaudissard *et al.*, 2017, Gauducheau *et al.*, 2017, Pietropaolo & Subashi, 2014).

Subjects of study 1 were adult (3 months old) C57BL/6J WT male mice, purchased from Janvier (Le Genest St Isle, France) and left undisturbed for two weeks before the beginning of behavioural testing. Subjects of studies 2, 3, 4 and 5 were adult (4-5 months old) male Fmr1-KO and their wild-type littermates (first generation model, (Dutch-Belgian Fragile X Consortium, 1994)), bred in our animal facility at Bordeaux University. C57BL/6JFmr1tm1Cgr/Nwu (B6) breeders were originally obtained from Neuromice.org (Northwestern University). Breeding trios were formed by mating two heterozygous Fmr1 females with a wild-type C57BL/6J male purchased from Janvier (Le Genest St Isle, France). After 2 weeks the sire was removed and the females were single caged and left undisturbed until weaning of the pups. Mice were weaned at 21 days of age and group-housed with their same-sex littermates (3–5/cage). On the same day, tail samples were collected for DNA extraction and subsequent PCR assessment of the genotypes as previously described (Dutch-Belgian Fragile X Consortium, 1994). Only litters including males of both genotypes (WT and KO) were used for experiments.

CD1 female mice (12±1 weeks old) purchased from Janvier (Le Genest St Isle, France) were used as social stimuli during the social interaction test for all studies. This strain has been selected for its high levels of sociability (Moles & D’amato F, 2000) and was previously employed in several social studies from our group on Fmr1-KO mice (Oddi *et al.*, 2015, Pietropaolo *et al.*, 2014, Pietropaolo *et al.*, 2011). Mice were group-housed (4-5/ cage) in standard cages (enriched with a cotton nestlet) and left undisturbed upon arrival two weeks before the social interaction test.

Subjects of study 6 were adult (4-5 months old) B6 Fmr1-KO2 males (the second generation Fmr1-KO mouse model, (Mientjes *et al.*, 2006)) and their wild-type littermates, bred in our animal facility at Bordeaux University, following the same procedures previously described for the Fmr1-KO model of the first generation. Mice were genotyped at weaning by tail PCR as described by Mientjes et al. (2006), and housed in unisexual groups in standard cages (enriched with a cotton nestlet) for the entire duration of the study.

All animals were group-housed in polycarbonate standard cages (33×15×14 cm in size; Tecniplast, Limonest, France), provided with litter (SAFE, Augy, France) and a stainless steel wired lid. Food (SAFE, Augy, France) and water were provided ad libitum. The animals were maintained in a temperature (22°C) and humidity (55%) controlled vivarium, under a 12:12 hr light–dark cycle (lights on at 7 a.m.). All experimental procedures were in accordance with the European Communities Council Directive of 24 November 1986 (86/609/EEC) and local French legislation.

### BEHAVIOURAL TESTS

To evaluate the therapeutic potential of CHLOR we have chosen behavioural tests that are currently used to evaluate FXS- and ASD-like behaviours. These include hyper-activity (the open field test), spatial memory (the Y maze test for spontaneous alternation), social deficits (direct social interaction test), repetitive behaviours (self-grooming test in a novel environment), and sensory hyper-responsiveness (acoustic startle test). The tests chosen have the following crucial advantages:

- they allow to detect behavioural abnormalities of *Fmr1*-KO mice that have been repeatedly and consistently described across several studies (for a review, see (Pietropaolo & Subashi, 2014))
- they allow a rapid (because of the short duration and the possibility of testing multiple animals at the same time) and mostly automatic assessment of mouse behavior
- they require minimal manipulation and stress of the subjects, thus allowing repeated testing of the same mice in multiple tests (and reducing the number of subjects needed).

All behavioral tests were carried out during the light phase of the cycle. Mice were habituated to the experimental room prior to all behavioral tests, being individually housed in standard polycarbonate cages provided with sawdust, food, and water bottles and left undisturbed for at least 10-15 min before testing began.

#### Open field test

this is one of the most widely used tests to assess locomotor activity and exploration in laboratory mice (Belzung *et al.*, 2005). The apparatus consisted of 4 white opaque plastic arenas (42×26×15cm). Each mouse was placed in the center of the arena and left free to explore it for 5 minutes. Automated Tracking of the videos obtained from a camera above the open field was performed with Ethovision (version 11, Noldus Technology, Wageningen, Netherlands) to analyze the total distance travelled.

#### Spatial memory in the Y maze test for spontaneous alternation

Spatial memory was assessed in a grey, plastic Y-maze, as previously described in details (Pietropaolo *et al.*, 2011). Briefly, during the first sample phase, access to the third novel arm was blocked by a door; mice were placed at the end of the start arm and allowed to freely explore the start and the other unblocked arm for 5 min before being returned to a waiting cage. After 2 min in the waiting cage, the test phase began: the door was removed; mice were placed at the end of the start arm and allowed to explore the entire maze for 2 min. Time spent in each arm of the maze was analyzed during both phases of the experiment as well as the distance travelled. Spontaneous alternation was assessed as follows: (Time spent in the novel arm/Time spent in all arms) x 100.

#### Direct social interaction test

Social interaction was assessed in a 30×15×14 cm cage, covered by a flat metal grid and with approximately 3 cm of sawdust on the floor, where male subjects were previously isolated for one hour. An unfamiliar CD1 female (3 months-old), was then introduced into the testing cage and left there for 3 min. Stimulus females were housed in unisexual groups in a female-only animal room and were in the non-estrous phase when tested (as assessed by the analysis of vaginal smears).

Testing sessions were recorded and videos analyzed with Observer XT (version 7, Noldus, The Netherlands). Time spent in affiliative behaviors of the male subject was evaluated, including sniffing the head and the snout of the partner, its anogenital region, or any other part of the body; allogrooming (grooming the partner) and traversing the partner’s body by crawling over/under from one side to the other.

#### Self-grooming test in a novel environment

The apparatus consisted of an unfamiliar Plexiglas cage (30×15×14 cm), covered by a flat metal grid and with approximately 3 cm of sawdust on the floor. A camera was placed in front of the cage for video recording. Mice were singly placed in the apparatus and allowed to explore for 20 minutes. Total time spent grooming was scored with Observer XT (version 7, Noldus, The Netherlands) by an observer who was blind to animals’ genotype and treatment.

#### Acoustic startle test

This test used a particular protocol involving low intensity stimuli that has been used so far only in Fmr1-KO mice (in both first and second generation models), where it has allowed detecting the sensory-hyper responsiveness of FX mice (Gaudissard *et al.*, 2017, Michalon *et al.*, 2014, Zhang *et al.*, 2014). The apparatus consisted of four acoustic startle chambers for mice (SR-LAB, San Diego Instruments, San Diego, CA, USA). Each comprised a nonrestrictive cylindrical enclosure made of clear Plexiglas attached horizontally on a mobile platform, which was in turn resting on a solid base inside a sound-attenuated isolation cubicle. A high-frequency loudspeaker mounted directly above the animal enclosure inside each cubicle produced a continuous background noise of 66 dBA and various acoustic stimuli in the form of white noise. Vibrations of the Plexiglas enclosure caused by the whole-body startle response of the animal were converted into analog signals by a piezoelectric unit attached to the platform. These signals were digitized and stored by a computer. The sensitivity of the stabilimeter was routinely calibrated to ensure consistency between chambers and across sessions.

A session began when the animals were placed into the Plexiglas enclosure. They were acclimatised to the apparatus for 5 minutes before the first trial began. Mice were then presented with pulses of white sound of 20 ms duration and varying intensity: +6, +12 +18 and +24 dB over background levels (namely 72, 78, 84 and 90 dB). Each intensity was presented 8 times, in a randomized order with variable intervals (10 sec to 20 sec) between the onset of each pulse. Mice were habituated to the boxes in the absence of acoustic stimuli 24hs prior to testing to reduce stress. A total of 130 readings of the whole-body startle response were taken at 0.5-ms intervals (i.e., spanning across 65 ms), starting at the onset of the pulse stimulus. The average amplitude (in mV) over the 65 ms was used to determine the stimulus reactivity and further averaged across trials.

### DRUG PREPARATION

All injectable solutions were freshly prepared on each experimental day. CHLOR (Chlorzoxazone, Sigma Aldrich, France), MET (Metformin, Sigma Aldrich, France) and GAB (Gaboxadol Hydrochloride, Sigma Aldrich, France) were dissolved in saline solution containing 1.25% DMSO (Sigma Aldrich, France) and 1.25% Tween80 (Sigma Aldrich, France). The same solution without drugs was used for the VEH control group. The doses of MET and GAB were chosen based on previous studies in Fmr1-KO mice (Gantois *et al.*, 2017, Olmos-Serrano *et al.*, 2011). In studies 1, 3, 4, 5 and 6, assessing acute effects, mice received a single i.p. injection of solutions containing either Veh or treatments (i.e., Chlor, MET or GAB, at the doses specified below) one hour before being submitted to the behavioural tests. In study 2, assessing chronic effects, mice received a daily i.p. injection of VEH solution, CHLOR (5mg/Kg) or MET (200mg/Kg) for 10 days, and behavioural tests were conducted 24hs after the last injection, in order to avoid acute effects.

### STATISTICAL ANALYSIS

Data were inspected for the identification of possible outliers using Grubbs’ test, or extreme studentized deviate ESD method. The number of outliers (1-2/group, if any) was in line with that seen in other similar studies conducted by this laboratory. Outliers were excluded from statistical analysis of the specific dataset and variable only; this explains the slight differences that may occur among tests in the number of animals per group. For each test, the exact N is indicated in the corresponding figure legend. Normality was assessed through the Shapiro-Wilks test for each experimental group and each variable of interest. Data from startle reactivity did not show a normal distribution at all stimulus intensities and were therefore subjected to natural logarithmic (ln) transformation in order to meet the normality requirements of ANOVA.

For all other variables, data distribution was found to be normal and all data were analyzed with ANOVA using treatment and genotype as the between-subject factors and adding stimulus intensity as the within-subject factor for the acoustic startle data. Post-hoc comparisons were performed when a significant interaction genotype x treatment was found using Tukey’s-Kramer test. Otherwise, separate one-way ANOVAs in each treatment group with genotype as the between subject factor were conducted, if appropriate. Data from the Y maze were analyzed using a one-sample t-test to evaluate the significant difference from the chance level, as done in previous studies (Oddi *et al.*, 2015, Vandesquille *et al.*, 2013). All analyses were carried out using Statview and PASW Statistics 18. Data are expressed as Mean ±SEM throughout the text.

## RESULTS

### Study 1: Dose-response study of the acute effects of CHLOR in WT mice

In our first study we have evaluated the behavioural effects of multiple doses of CHLOR in WT mice of the same background (B6) used for our Fmr1-KO mice. The aim of this study was to identify the optimal dose to be used in the subsequent experiments, that is, the highest dose having no effect in WT mice. The following doses were used: 5, 10 and 20 mg/Kg that were administered to different groups of mice 1hr before the open field followed (after a 5 min interval) by the acoustic startle test. The results showed that CHLOR treatment clearly affected the locomotor activity of WT mice in the open field (Fig.1-A), starting from the dose of 10 mg/Kg [treatment effect: F(3,34)=6.12, p<0.01; post-hoc: D10 and D20 versus Veh, D10 versus D5, p<0.05]. No effects of the treatments were observed on acoustic startle response, where only a significant effect of stimulus intensity was found, as expected (Fig.1-B).

**Fig. 1:**
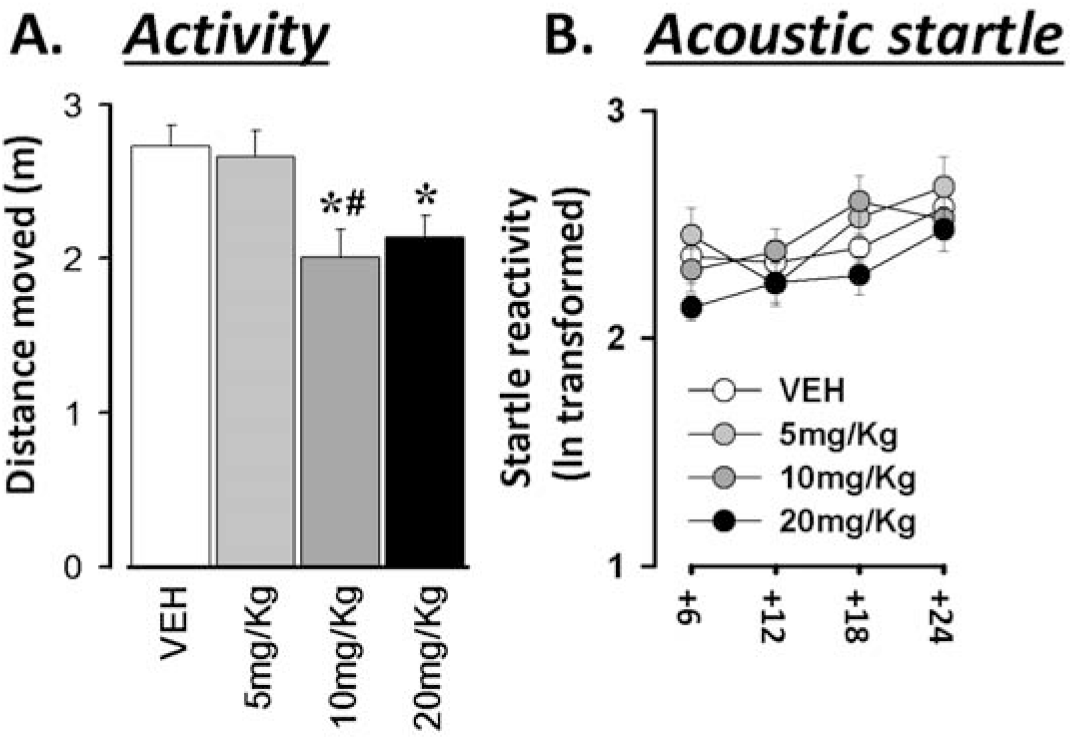
Locomotion in the open field (A) and acoustic startle response (B) in a dose-response study in WT mice: n=9-10; *=p<0.05, versus VEH; # versus D5. Data are expressed as Mean ±SEM. Stimulus intensities are expressed over (+) a background of 66dB.

### Study 2: Comparing CHLOR versus Metformin: Chronic effects in Fmr1-KO mice

As expected, Fmr1-KO mice treated with VEH showed hyperactivity (Fig.2-A), and reduced social interaction (Fig.2-B); these abnormalities were not attenuated by acute MET administration, but were eliminated by CHLOR [interaction genotype x treatment, respectively: F(2,34)=7.05, and F(2,30)=5.32, p<0.01 and =0.01; post-hoc: KO-VEH versus WT-VEH, KO-CHLOR versus KO-VEH: p<0.05; for hyperactivity, also KO-MET versus WT-MET and KO-CHLOR: p<0.05]. Similarly, only CHLOR eliminated the exaggerated acoustic response of KO mice that was especially marked at the highest stimulus intensities; nonetheless, this acoustic abnormality seemed slightly less pronounced following MET treatment [Fig.2-D; interaction genotype x intensity: F(3,102)=9.41, p<0.0001; interaction genotype x treatment F(2,34)=4.48, p<0.05; genotype effects in separate ANOVAs: in VEH F(1,11)=10.03, p<0.01; in MET F(1,11)=7.13, p<0.05, in CHLOR, n.s.]. KO-VEH mice also spent more time performing self-grooming (Fig.2-C), that is considered an index of anxiety/repetitive behavior, and this abnormal phenotype was eliminated by both MET and CHLOR chronic administration [interaction genotype x treatment: F(2,30)=8.24, p<0.01; post-hoc: KO-VEH versus WT-VEH, KO-CHLOR and KO-MET: p<0.05].

**Fig. 2:**
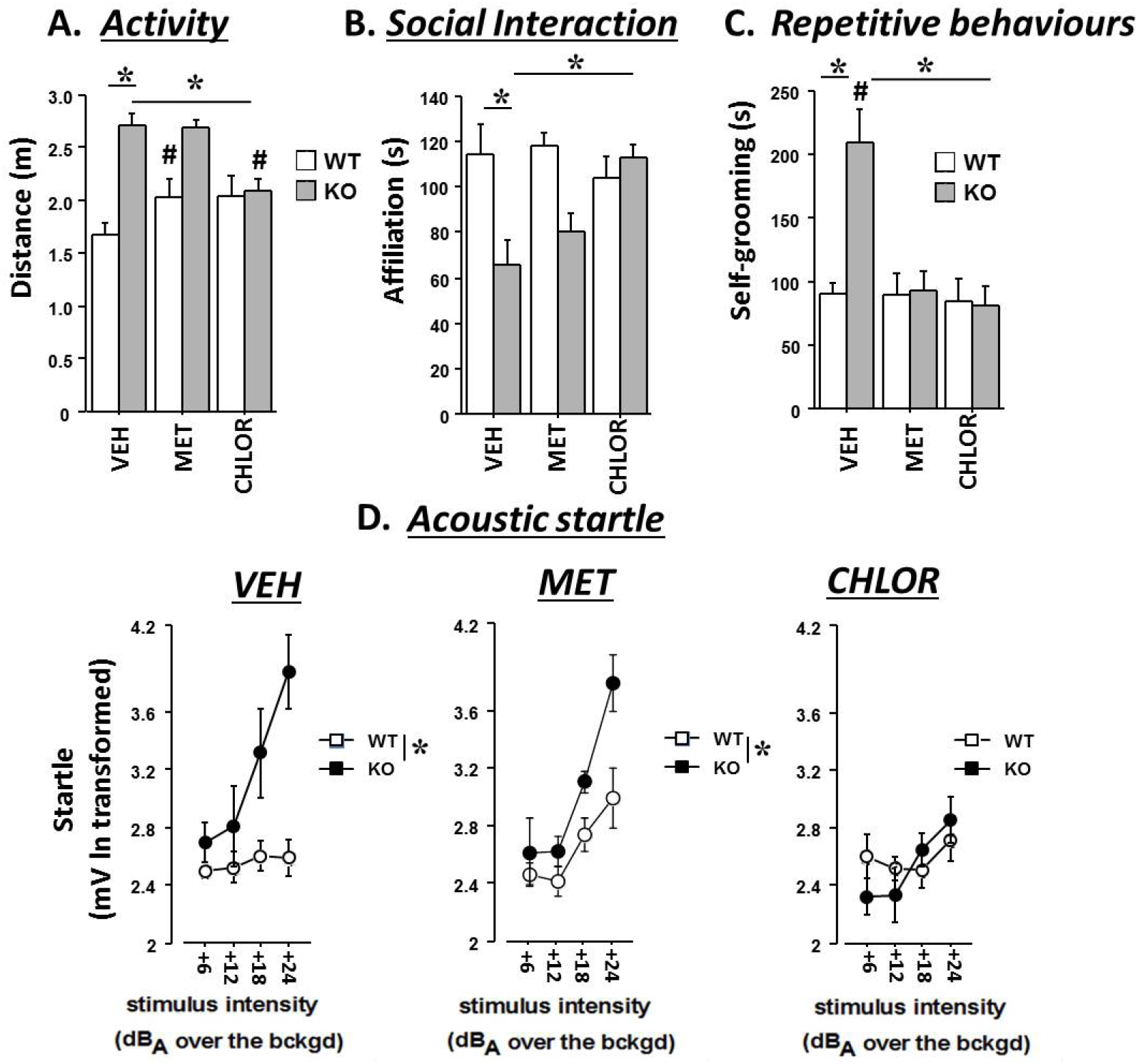
Chronic effects of chlorzoxazone (CHLOR; 5mg/Kg) versus Metformin (MET, 200mg/Kg): Behavioural tests were conducted after 10 days of i.p. injections (with a 24hs interval from the last injection) and they included open field test for locomotor activity (A), direct social interaction with a WT female (B), evaluation of self-grooming in a novel cage (C) and acoustic startle response test (D). n=6-8; *=p<0.05; #= versus KO-MET. Stimulus intensities are expressed over (+) a background of 66dB.

To include potential non-behavioural effects (Fig.3), we investigated whether chronic CHLOR or MET could rescue the macro-orchidism known to affect Fmr1-KO mice (as FXS patients). None of the treatments eliminated the enhanced testicular weight of Fmr1-KO mice [Fig.3-A, genotype effect F(1,34)=19.51, p<0.0001; all other effects and interaction: n.s.]. Body weight was slightly reduced after both CHLOR and MET chronic treatments, an effect that was equally observed in WT and KO mice [Fig.3-B, treatment effect on body weight loss: F(2,34)=10.25, p<0.001; post-hoc: CHLOR and MET versus VEH: p<0.05].

**Fig. 3:**
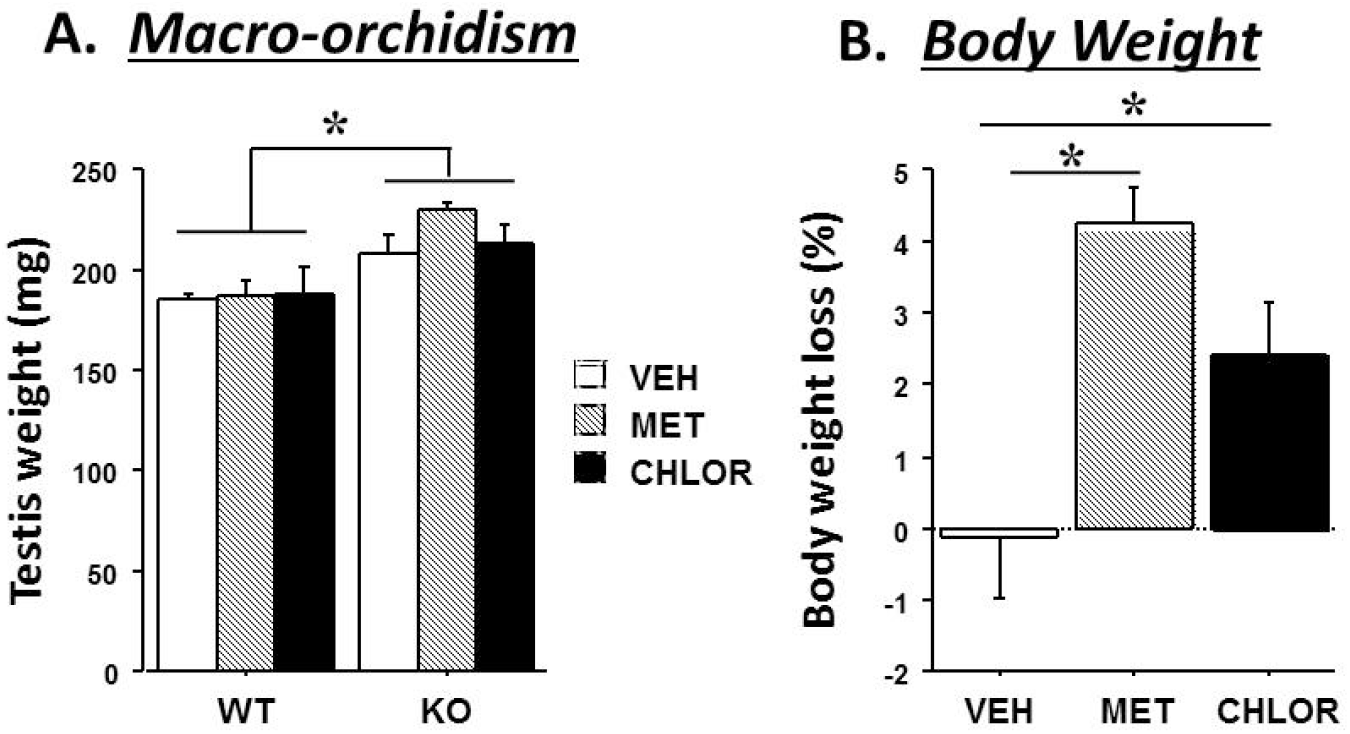
Chronic non-behavioural effects of CHLOR (5mg/Kg) and MET (200mg/Kg): Testis weight (A) and body weight (B; averaged across genotypes) were assessed after 10 days of i.p. injections. n=6-8; *=p<0.05.

### Study 3: Comparing CHLOR versus Metformin: Acute effects in Fmr1-KO mice

Mice received a single i.p. injection of Veh solution, Chlor (5mg/Kg) or MET (200mg/Kg) and were submitted one hour afterwards to the open field test, social interaction test, Y maze test for spontaneous alternation and acoustic startle test.

As expected, Fmr1-KO mice treated with VEH showed hyperactivity (Fig.4-A), and reduced social interaction (Fig.4-B); these abnormalities were not attenuated by acute MET administration, but were eliminated by CHLOR [interaction genotype x treatment, respectively: F(2,53)=6.13, and F(2,48)=8.85, both p<0.01; post-hoc: KO-VEH versus WT-VEH, KO-CHLOR versus KO-VEH and KO-MET: p<0.05; for hyperactivity, also WT-MET versus KO-MET: p<0.05]. CHLOR also restored spatial memory in the Y maze in Fmr1-KO mice (t-test, p<0.05, Fig.4-C), while KOs treated with MET showed a performance similar to those receiving VEH, i.e., not different from the chance level (t-test, n.s., Fig.4-C). Finally, Fmr1-KO mice treated with VEH displayed enhanced acoustic startle, especially at the highest stimulus intensities (Fig. 4-D), and this effect was eliminated by acute CHLOR, but not MET [overall effects of genotype: F(1,53)=7.69 p<0.01 and treatment F(2,53)=3.33, p<0.05; interaction genotype x treatment x stimulus intensity: F(6,159)=2.09, p=0.05; genotype effects in separate ANOVAs: in VEH F(1,16)=6.06, p<0.05; in MET F(1,19)=5.66, p<0.05, in CHLOR, n.s.].

**Fig. 4:**
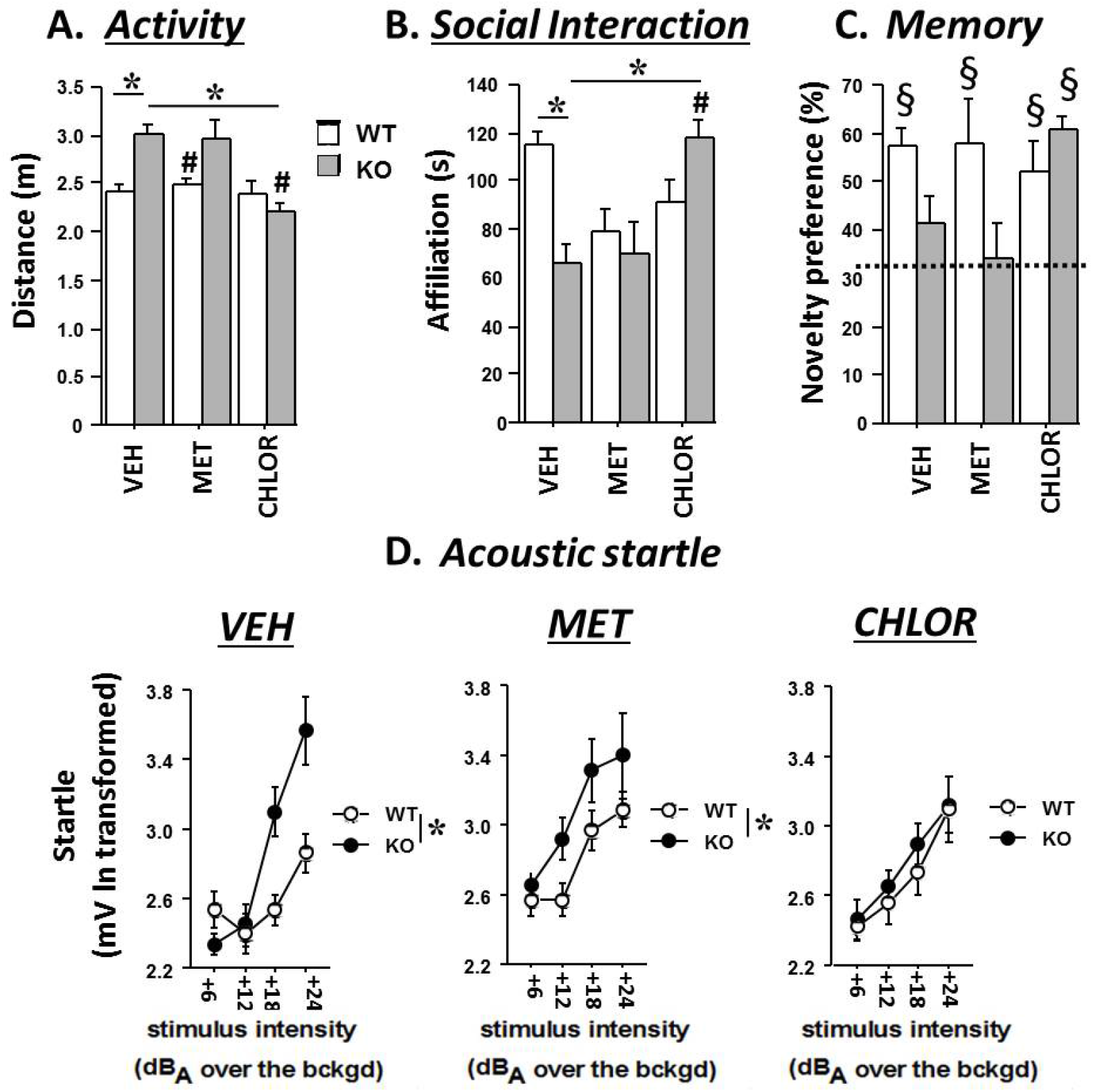
Acute effects of chlorzoxazone (CHLOR; 5mg/Kg) versus Metformin (MET, 200mg/Kg). Behavioural tests were conducted 1h after a single i.p. injection and they included open field test for locomotor activity (A), direct social interaction with a WT female (B), Y maze test for spatial memory (C) and acoustic startle response test (D). n=8-10; *=p<0.05; #= versus KO-MET, §=versus chance level. Stimulus intensities are expressed over (+) a background of 66dB

### Study 4: Comparing CHLOR versus Gaboxadol: Acute effects in Fmr1-KO mice

Mice received a single i.p. injection of Veh solution, Chlor (5mg/Kg) or GAB (either at the dose of 0.5 or 3 mg/Kg). One hour after injection, animals were submitted to the open field test, social interaction test, and acoustic startle test.

Once again, Fmr1-KO mice treated with VEH showed hyperactivity (Fig.5-A), reduced social interaction (Fig.5-B) and enhanced acoustic startle, especially at the highest stimulus intensities (Fig. 5-C); all these abnormalities were not significantly attenuated by acute GAB administration at either doses, but were instead eliminated by CHLOR [interaction genotype x treatment on total distance and affiliation time, respectively: F(3,62)=4.91, and F(3,64)=6.26, both p<0.01; post-hoc: KO-VEH versus WT-VEH, KO-CHLOR versus KO-VEH: p<0.05; for startle data, interaction genotype x treatment x stimulus intensity: F(3,192)=3.94, p<0.001; post-hoc: KO-VEH versus WT-VEH, KO-CHLOR versus KO-VEH: p<0.05].

**Fig. 5:**
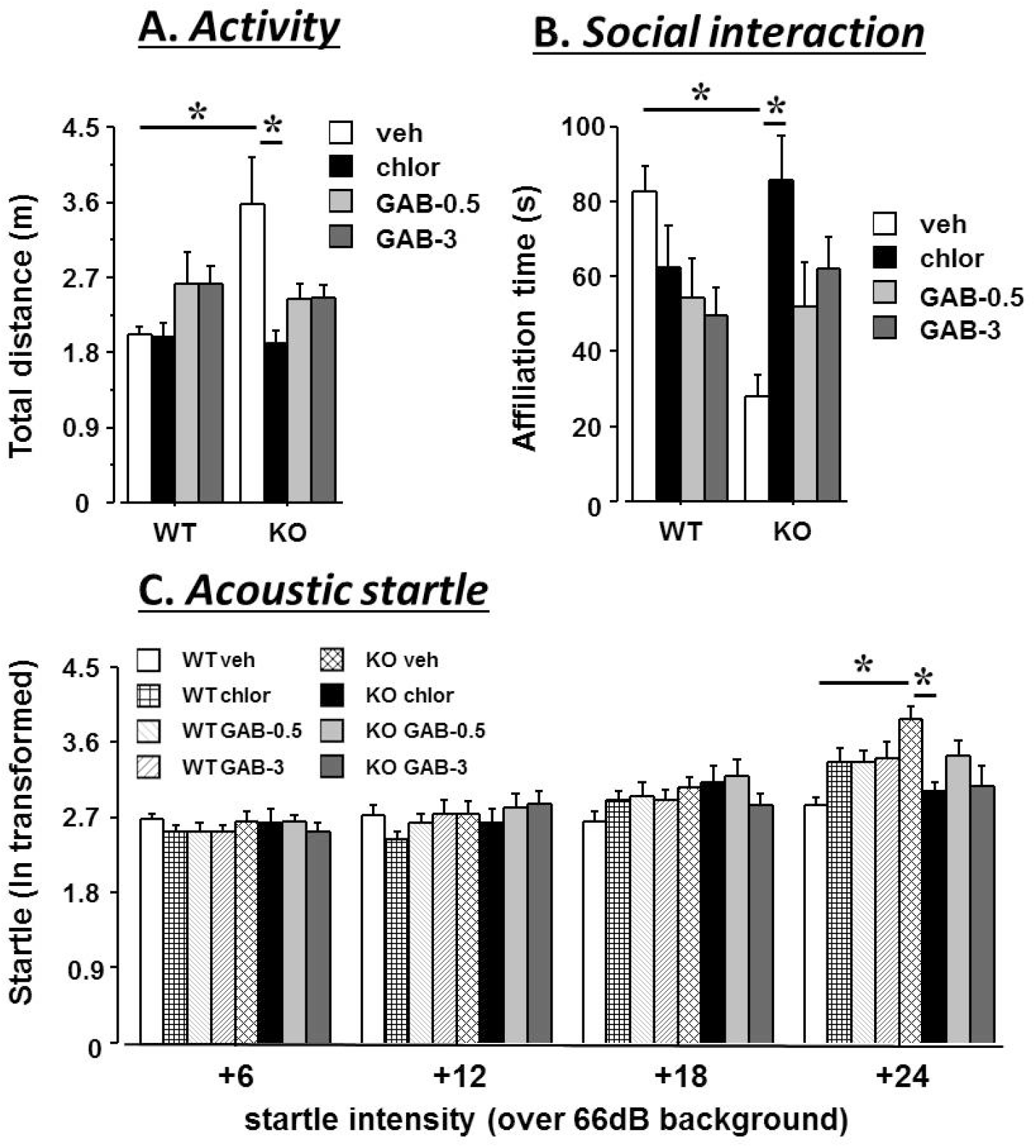
Acute effects of CHLOR and two doses of Gaboxadol (GAB) in the open field (A), social interaction (B) and acoustic startle (C) tests. n=9-10; *=p<0.05. Doses are expressed as mg/Kg; treatments were administered 1 hr before testing. Stimulus intensities are expressed over (+) a background of 66dB.

### Study 5: Dose-response of Chlor acute effects in Fmr1-KO mice

We then performed a dose-response study on the acute effects of CHLOR in Fmr1-KO mice. The aim of this study was to evaluate whether a dose lower than the one used so far, i.e., 5 mg/Kg, could be sufficient to induce acute beneficial behavioural effects in Fmr1-KOs. The following doses of CHLOR were used: 1.25, 2.5 3.75, and 5 mg/Kg corresponding to 25, 50, 75 and 100% of the original dose. A single i.p. injection of either the Vehicle or one of the CHLOR solutions was administered in Fmr1-KO mice and their WT littermates 1 hour before the beginning of behavioral testing. Hyperactivity in the open field and acoustic startle response were assessed as in previous studies.

A dose effect of CHLOR was found on hyperactivity of KO mice, the dose of 5 mg/Kg being the most effective in reducing the hyper-locomotion of mutant animals, followed by the 3.75mg/Kg dose [Fig. 6A: interaction genotype x treatment on the distance moved: F(4,68)= 10.08, p<0.0001; post-hoc: KO-VEH versus WT-VEH and versus KO-CHLOR5 and KO-CHLOR3.75; KO-CHOR1.25 and KO-CHLOR2.5 versus WT with same treatments: p<0.05]. Similar results were observed on the acoustic startle, but only the highest dose of 5 mg/Kg was able to fully rescue the acoustic hyper-responsiveness of KO mice [Fig. 6B: interaction genotype x treatment: F(4,68)= 2.26, p=0.07; interaction genotype x treatment x stimulus intensity: F(12,204)=1.67, p=0.07; genotype effects in separate ANOVAs: in VEH F(1,14)=3.56, p<0.01; in CHLOR1.25 F(1,14)=4.75, p<0.05, in CHLOR2.5 F(1,14)=8.44, p<0.05, in CHLOR3.75 F(1,13)=9.14, p<0.05, in CHLOR5 n.s.].

**Fig. 6:**
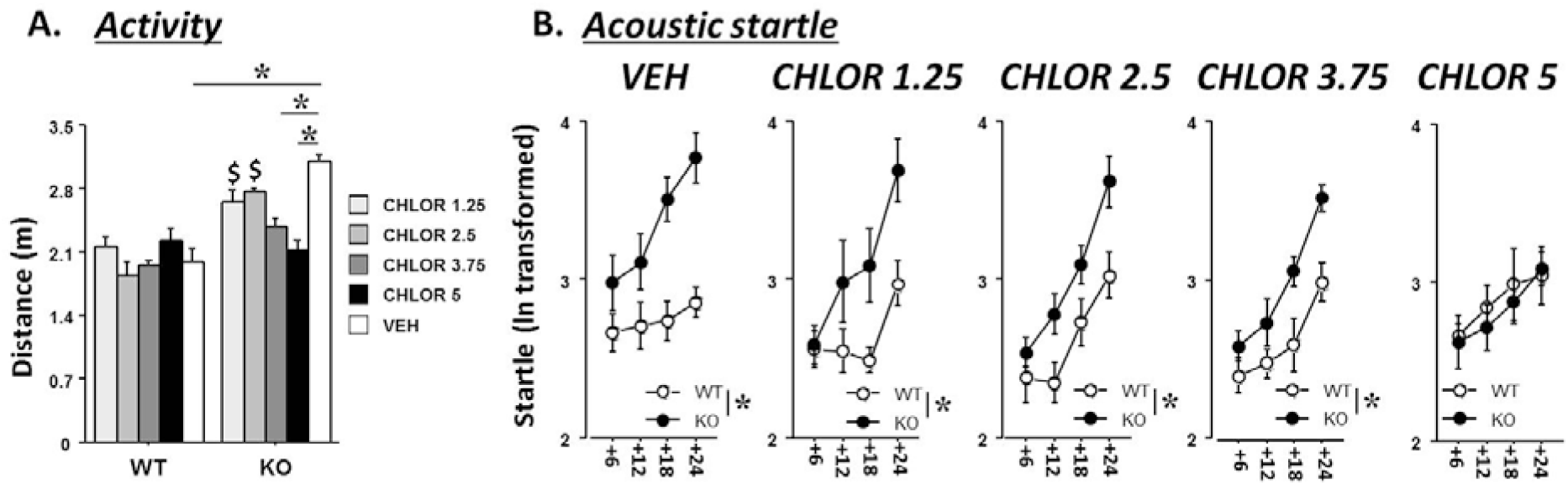
Dose-response study on the acute effects of CHLOR in the open field (A) and acoustic startle (B) tests. n=9-10; *=p<0.05. $ versus WT with same treatments: p<0.05. Doses are expressed as mg/Kg; treatments were administered 1 hr before testing. Stimulus intensities are expressed over (+) a background of 66dB.

### Study 6: Acute effects of Chlor in the Fmr1-KO mouse model of second generation (Fmr1-KO2 mouse line)

The Fmr1-KO mouse model we have used so far was engineered in 1994 (The Dutch-Belgian Fragile X Consortium, 1994) and is still the most widely used in the research on FXS and ASD for behavioural, molecular and electrophysiological studies. Nonetheless, the Fmr1-KO mouse line of the second generation can also be used, differing from the original Fmr1 model because of the full absence not only of the FMRP protein, but also of related mRNAs (Mientjes *et al.*, 2006). This model, known as Fmr1-KO2, is less widely used in neurobiological research and mostly for electrophysiological and molecular investigations (Zhang *et al.*, 2014). Indeed, when we have performed an extensive behavioural characterization of Fmr1-KO2 mice at adulthood, we detected a less varied ASD- and FXS-like phenotype compared to the first Fmr1-KO model (Gaudissard *et al.*, 2017). For example, we did not observe hyperactivity and social deficits in KO2 mice (Gaudissard *et al.*, 2017), while exaggerated acoustic startle response was confirmed as a robust behavioural alteration also of these mutants (Gaudissard *et al.*, 2017, Zhang *et al.*, 2014). Hence, we assessed here whether a single injection of the dose of 5mg/Kg of CHLOR could rescue the enhanced acoustic startle of Fmr1-KO2 mice tested one hour afterwards. We indeed applied exactly the same experimental procedures previously used for Fmr1-KOs to Fmr1-KO2 mice (Gaudissard *et al.*, 2017).

The results confirmed that acute CHLOR eliminated the exaggerated startle response of KO2 mice (Fig. 7), as previously demonstrated in the KO model of the first generation [overall effects of genotype: F(1,24)=16.07, p<0.001 and treatment F(1,24)=6.14, p<0.05; interaction genotype x treatment: F(3,72)=4.45, p<0.05; genotype effects in separate ANOVAs: in VEH F(1,11)=38.71, p<0.0001; in CHLOR, n.s.].

**Fig. 7:**
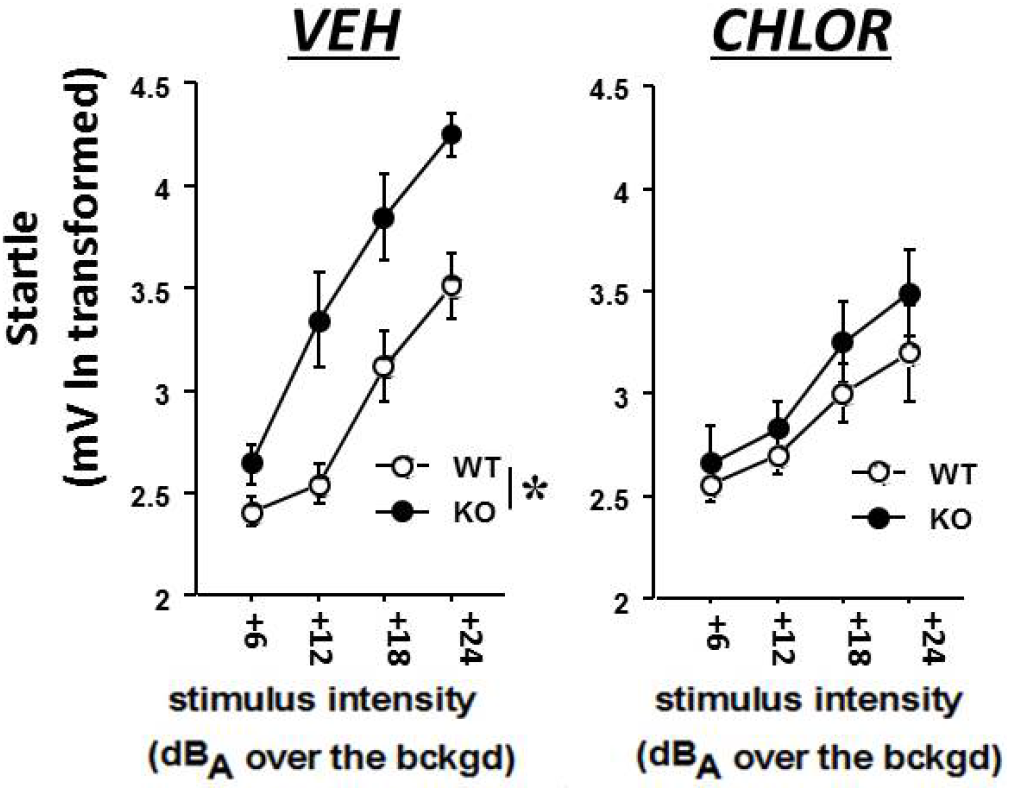
Acute effects of CHLOR (5 mg/Kg) on the acoustic startle response in Fmr1-KO2 mice, i.e., the second generation mouse model of FXS: n=6-8; *=p<0.05. Treatments were administered 1h before testing. Stimulus intensities are expressed over (+) a background of 66dB.

## DISCUSSION

Our data demonstrate for the first time the efficacy of Chlorzoxazone in rescuing all the FXS-like behavioural abnormalities of Fmr1-KO mice (using models of both first and second generation). The therapeutic impact of chlorzoxazone (CHLOR) was demonstrated both in acute and chronic administration, showing higher efficacy than other two molecules proposed to treat FXS, i.e., Metformin (MET) and Gaboxadol (GAB). Furthermore, no aversive side effects were observed following chronic (10 days) treatments with CHLOR, as only a slight weight loss (2% of initial body weight) was detected (Metformin had instead a pronounced effect on body weight, i.e., inducing a weight loss of about 4%). Overall, the dose of 5 mg/Kg seemed the most appropriate to induce the most varied and robust behavioural rescue of the FXS-phenotype of Fmr1-KO mice, a dose that is quite in line with the concentration range used in human posology (i.e., 250-500 mg/day for an adult subject (Hohmann *et al.*, 2019)). Nonetheless, acute therapeutic effects of a lower dose (corresponding to 75% of the original dose of 5 mg/Kg) were found here on hyperactivity, thus suggesting that a reduction in the daily dose may be possible for clinical applications. This is an important issue, as using microdoses of CHLOR may further limit its potential toxicity (Hohmann *et al.*, 2019), a problem that may be especially relevant because of the long duration of administration necessary for young patients. Interestingly, the effects of CHLOR were specific for the behavioural FXS-like phenotypes, as no effects on macro-orchidism were detected, suggesting that brain rather than peripheral BK channels may be preferentially targeted by our treatment. This issue may be further investigated through electrophysiological studies assessing the effects of CHLOR on BKCa functionality in the brain of Fmr1-KO mice, confirming its efficacy in rescuing the deficits observed in these mutants.

Our results support the view that molecules acting as BKCa agonists may be a valid therapeutic strategy for FXS, i.e., an idea that we have already proposed in previous studies in Fmr1-KO mice, where we demonstrated that the acute administration of BMS, another BKCa channel opener, was able to eliminate several behavioural abnormalities in FXS mice (Carreno-Munoz *et al.*, 2018, Hebert *et al.*, 2014, Zhang *et al.*, 2014), as well as their hippocampal dendritic alterations (Hebert *et al.*, 2014) and cortical hyper-excitability (Zhang *et al.*, 2014). Nonetheless, CHLOR seems a more valuable therapeutic agent for clinical application than BMS, as the latter suffers from limited efficacy in chronic protocols of administration and short half-life (Hebert *et al.*, 2014), i.e., two problems that do not affect CHLOR (Hohmann *et al.*, 2019, Witt *et al.*, 2016). The comparison between CHLOR and other two treatments recently proposed for FXS further strengthens the value of this molecule as a treatment for FXS; here, we could not detect any acute behavioural effect of MET, in agreement with previous reports on the Fmr1-KO model (Gantois *et al.*, 2017), and we found limited chronic effects of MET, i.e., on repetitive behaviours and, more slightly, on social deficits (Fig.2-B and –C), that were largely less marked than those induced by chronic CHLOR. For GAB, slight acute effects were detected in a dose-dependent manner, on hyperactivity and social interaction (Fig.5-A and B), in line with previous results (Olmos-Serrano *et al.*, 2011), but again, CHLOR was more powerful in inducing a full behavioural rescue of Fmr1-KO mice.

In conclusion, our findings show the relevance of BKCa as a therapeutic target for neurological disorders, as well as of CHLOR as a molecule of high interest for clinical applications. Recently, CHLOR has been shown exerting beneficial effects on the cognitive deficits and amyloid pathology in a mouse model of Alzheimer disease, by rescuing alterations in BKCa functionality and inflammatory imbalances (Bai & Ma, 2020). It is therefore possible that BKCa channels may represent a therapeutic target common to multiple disorders of the central nervous system (Bailey *et al.*, 2019). This may be particularly interesting for neurodevelopmental disorders, as alterations in BKCa expression (in particular of the KCNMA1 major channel subunit) and functionality have been demonstrated in several of these pathologies (Bailey *et al.*, 2019), including FXS and ASDs. The link between these two syndromes is largely supported (Hatton *et al.*, 2006), as they are intertwined at the molecular (Darnell *et al.*, 2011, Parikshak *et al.*, 2013) and behavioural level (Bailey *et al.*, 2001, Bailey *et al.*, 1998, Brock & Hatton, 2010, Rogers *et al.*, 2001): indeed, all FXS patients meet some aspects of the DSM criteria for ASD and at least 50% meet the full criteria (Roberts *et al.*, 2007). Since for ASD no universally accepted animal model exists, there is a growing interest in using FXS mouse models to study this complex category of developmental pathologies (Bernardet & Crusio, 2006, Oddi *et al.*, 2013, Pietropaolo & Subashi, 2014). Thus, identifying novel therapeutic strategies for FXS may critically contribute to advance our understanding of the etiopathological pathways common to multiple developmental and neurological disorders.

## ACKNOWLEDGEMENTS

The authors thank Delphine Gonzales and the genotyping facility of Neurocentre Magendie, funded by Inserm and LabEX BRAIN ANR-10-LABEX-43, for animal genotyping. We thank Elodie Poinama and Renata Hermez for their expert animal care, Thierry Lafont for technical assistance, Christophe Halgand and Loic Grattier for informatics support.

